# Tethering NDC80-NUF2 to microparticles is sufficient to enable their biorientation in the spindle

**DOI:** 10.1101/2024.03.09.582646

**Authors:** Kohei Asai, Yuanzhuo Zhou, Tomoya S. Kitajima

**Affiliations:** Laboratory for Chromosome Segregation, RIKEN Center for Biosystems Dynamics Research (BDR), Kobe, Japan; Graduate School of Biostudies, Kyoto University, Kyoto, Japan

**Keywords:** Chromosome segregation, biorientation, kinetochore, oocyte, live imaging

## Abstract

Faithful chromosome segregation requires biorientation, where the pair of kinetochores on the chromosome establish bipolar microtubule attachment. The integrity of the kinetochore, a macromolecular complex built on centromeric DNA, is required for biorientation, but components sufficient for biorientation remain unknown. In this study, we show that tethering the outer kinetochore heterodimer NDC80-NUF2 to the surface of microbeads with no bipolar cue is sufficient for them to establish a biorientation-like state in mouse oocytes. NDC80-NUF2 microbeads efficiently and stably align at the spindle equator, forming bipolar microtubule attachments. Furthermore, they can self-correct alignment errors. They align independently of the outer kinetochore proteins SPC24-SPC25, KNL1, the MIS12 complex, or inner kinetochore proteins. Aurora-mediated NDC80 phosphoregulation promoted microbead alignment under challenging conditions. Interestingly, larger microbeads align more rapidly, suggesting that large platform size enhances NDC80-NUF2-mediated biorientation. This study shows a biohybrid kinetochore design for synthetic biorientation of microscale particles in cells.

## Introduction

Accurate chromosome segregation during cell division is essential for genome stability. Central to this process is chromosome biorientation, wherein microtubules attach to the pair of kinetochores on the chromosome from the opposite spindle poles. Kinetochores, complex macromolecular assemblies that form on centromeric DNA, are composed of over 100 different proteins in vertebrates ^1,2^. These kinetochore and centromere proteins can be categorized into at least three groups: (1) outer kinetochore proteins that form connections with spindle microtubules, (2) inner kinetochore proteins that mediate the linkage of outer kinetochore proteins with centromeric DNA, and (3) inner centromere proteins that localize between the kinetochore pair. Biorientation requires the selective stabilization of bipolar kinetochore-microtubule attachment, which is thought to be achieved by the coordinated actions of these kinetochore and centromere proteins. One model suggests that biorientation is mediated by spatial separation of kinetochore-microtubule attachment sites from the inner centromere, when microtubules pull the kinetochore pair towards opposite poles and induce intra-kinetochore stretching ^3–8^. This model is supported by substantial evidence that phosphorylation mediated by Aurora, an inner centromere kinase required for biorientation, reduces the microtubule-binding activity of outer kinetochore proteins such as the NDC80 complex ^9–11^. However, the inner centromere localization of Aurora kinase is not strictly required for accurate chromosome segregation ^12–14^, suggesting the existence of additional mechanisms for biorientation. An alternative, Aurora-independent model suggests that kinetochores possess an intrinsic capacity to stabilize microtubule attachment in response to tension ^15,16^. In support of this model, purified yeast kinetochores containing nearly the complete set of both outer and inner kinetochore proteins, but lacking inner centromere proteins such as Aurora, can directly stabilize microtubule attachment when subjected to tension *in vitro* ^15^. This tension-dependent stabilization relies on the recruitment of Stu2 (known as chTOG in mammals) to the outer kinetochore by the NDC80 complex ^16,17^. However, it remains unclear whether the NDC80 branch of the outer kinetochore alone is sufficient to establish biorientation in cells.

## Results

### NDC80-NUF2 recruitment enables microbeads to align

The NDC80-NUF2 heterodimer is an outer kinetochore subcomplex that contains microtubule-binding domains at their N-termini (Fig. 1a). It serves as a major anchor for load-bearing kinetochore-microtubule attachment ^10,11,18,19^. We hypothesized that NDC80-NUF2 recapitulates the microtubule-binding properties of kinetochores when bound to the surface of microbeads. Mouse oocytes were chosen as a model living system for functional assays because microbeads can be microinjected into them at the prophase-arrested stage, which can be released synchronously into M-phase (meiosis I). We microinjected mRNAs encoding NDC80 with a GFP tag at its C-terminus (NDC80-GFP) and NUF2, and then microinjected anti-GFP-conjugated microbeads (1.7–2.6 µm in diameter) into the cytoplasmic region near the nucleus of prophase-arrested oocytes (Fig. 1a, Supplementary Video 1). After one hour of incubation, entry into M-phase was induced. At metaphase (5 hours after nuclear envelope breakdown, NEBD), we observed that NDC80-GFP-coated microbeads (hereafter referred to as NDC80-NUF2 microbeads, see below) were located in close proximity to chromosomes, whereas control GFP-coated microbeads were located away from chromosomes, presumably outside the spindle (Fig. S1a). These data suggested that NDC80-NUF2 microbeads interact with spindle microtubules, which was consistent with our expectation. Surprisingly, however, 3D inspection of microbead positions revealed that NDC80-NUF2 microbeads were well aligned at the metaphase plate (Fig. S1a). This observation suggests that NDC80-NUF2 recruitment enables microbeads to align at the metaphase plate.

**Fig. 1:**
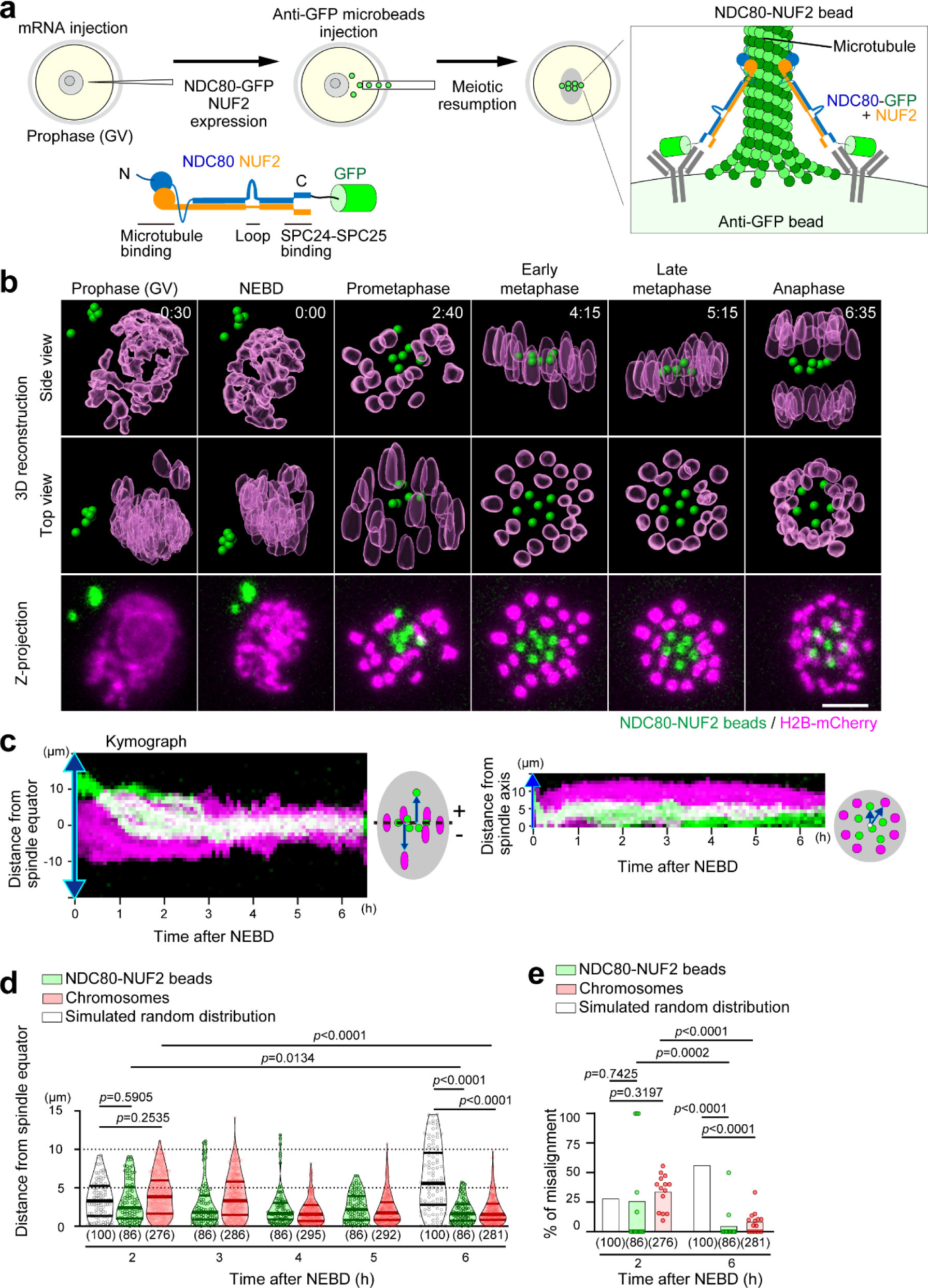
NDC80-NUF2 suffices microbead alignment. **(a)** Experimental design. NDC80-NUF2 beads were constructed through the coexpression of NDC80-GFP and NUF2 via mRNA microinjection, followed by anti-GFP bead microinjection, into the cytoplasm of prophase-arrested, germinal vesicle (GV) stage mouse oocytes. Bead dynamics were monitored following the induction of meiotic resumption. **(b)** Live imaging. 3D-reconstructed and z-projection images of chromosomes (H2B-mCherry, magenta) and NDC80-NUF2 beads (green). Time after NEBD (h:mm). Scale bar, 10 µm. **(c)** Kymograph. Images in **(b)** were kymographed with projected signals based on the spindle axis. The vertical axis of the kymographs shows the distance from the spindle equator (left) and the distance from the spindle axis (right), respectively. **(d)** Alignment efficiency. The distances of NDC80-NUF2 beads and chromosomes from the spindle equator were quantified. Simulated random distribution of particles within a spindle-like ellipsoid (20 µm at 2 h or 30 µm at 6 h in length and 20 µm in width) is used as a reference. The number of beads or chromosomes (from 14 oocytes in 3 independent experiments) is indicated in parentheses. P-values were calculated with Kruskal-Wallis test with Dunn’s correction. **(e)** Misalignment frequency. Fractions of misaligned NDC80-NUF2 beads and chromosomes in **(d)** are shown. “Misaligned” refers to positions over 5 µm from the spindle equator. Each dot shows the data of an oocyte. P-values were calculated with Fisher’s exact test.

We investigated whether the microbead alignment is mediated by NDC80-NUF2-specific properties. First, we tested other outer kinetochore components. KNL1 is an outer kinetochore protein with a microtubule-binding domain at its N-terminus ^10,20^. In contrast to NDC80-NUF2 microbeads, microbeads binding to C-terminally GFP-tagged KNL1 (KNL1-GFP) did not align in the spindle (Fig. S1a). Similarly, microbeads binding to MIS12-GFP, a component of the outer kinetochore MIS12 complex ^10^, did not show alignment (Fig. S1a). Second, we deleted or mutated the loop domain of NDC80, which is critical for the NDC80-NUF2 function (Fig. S1b) ^21–23^. Neither the loop-deleted nor the loop-mutated NDC80-NUF2 aligned microbeads, indicating that the intact loop is required and the N-terminal microtubule-binding domain is not sufficient for microbead alignment. Finally, expression of NDC80-GFP alone without NUF2 did not result in microbead alignment (Fig. S1c), suggesting that NDC80-NUF2 heterodimerization, likely facilitated by exogenous supply of NUF2, is critical for alignment ability. These results suggest that NDC80-NUF2 heterodimer-specific properties enable microbead alignment.

### Efficient alignment

We investigated how NDC80-NUF2 microbeads align using confocal live imaging. Analysis of microbead dynamics in 3D showed that NDC80-NUF2 microbeads gradually aligned with kinetics similar to chromosomes (Fig. 1b–e, Supplementary Video 2). Kymograph analysis along the spindle axis showed that the alignment of NDC80-NUF2 microbeads was stably maintained until the onset of anaphase, as observed for chromosomes (Fig. 1c). Interestingly, NDC80-NUF2 microbeads occupied the inner region of the metaphase plate, while chromosomes were ejected to the outer region (Fig. 1c, Fig. S1d). The microbeads were aligned at regular intervals similar to chromosomes (Fig. S1e). These results demonstrate that NDC80-NUF2 microbeads establish stable alignment with a preference for positioning at the inner region of the metaphase plate.

### Robust alignment

Metaphase is occasionally accompanied by one or a few misaligned chromosomes, which are normally corrected to align by the onset of anaphase. Similarly, we occasionally observed one or two misaligned NDC80-NUF2 microbeads in early metaphase (Fig. S2a). Tracking these microbeads in 3D showed their transient poleward movement, which was subsequently corrected to movement toward the metaphase plate, resulting in recovery of alignment (Fig. S2a–c, Supplementary Video 3). Consequently, at 5 minutes prior to anaphase onset, all of the NDC80-NUF2 microbeads we observed were aligned (n = 84 in 14 oocytes), positioning within 5 μm from the spindle equator (Fig. S2d). Thus, NDC80-NUF2 microbeads are capable of correcting their own misalignment, similar to chromosomes.

### Establishment of a biorientation-like state with bipolar microtubule attachment

These observations strongly suggest that NDC80-NUF2 microbeads are capable of establishing bipolar microtubule attachment. We examined microtubule attachment using HURP, a kinetochore fiber marker ^24^. At metaphase, when microbeads and chromosomes were fully aligned, the vast majority of NDC80-NUF2 microbeads (34/37, 92%) showed bipolar attachment to the end of HURP-decorated microtubule bundles, similar to the kinetochore pair of chromosomes (Fig. 2a, b, “Metaphase”, Supplementary Video 4). In contrast, at prometaphase, when alignment was not completed, bipolar attachment of HURP-decorated microtubule bundles was less frequent (Fig. 2a, b, “Prometaphase”). Notably, at this stage, aligned microbeads were more frequently attached to bipolar HURP-decorated microtubule bundles compared to unaligned microbeads (Fig. 2a, c). These features were similar to the kinetochore pair of chromosomes (Fig. 2a–c). Thus, alignment of NDC80-NUF2 microbeads is associated with bipolar end-on attachment to microtubule fibers. Based on these results and the fact that they exhibit stable alignment, we conclude that NDC80-NUF2 microbeads are capable of establishing a biorientation-like state.

**Fig. 2:**
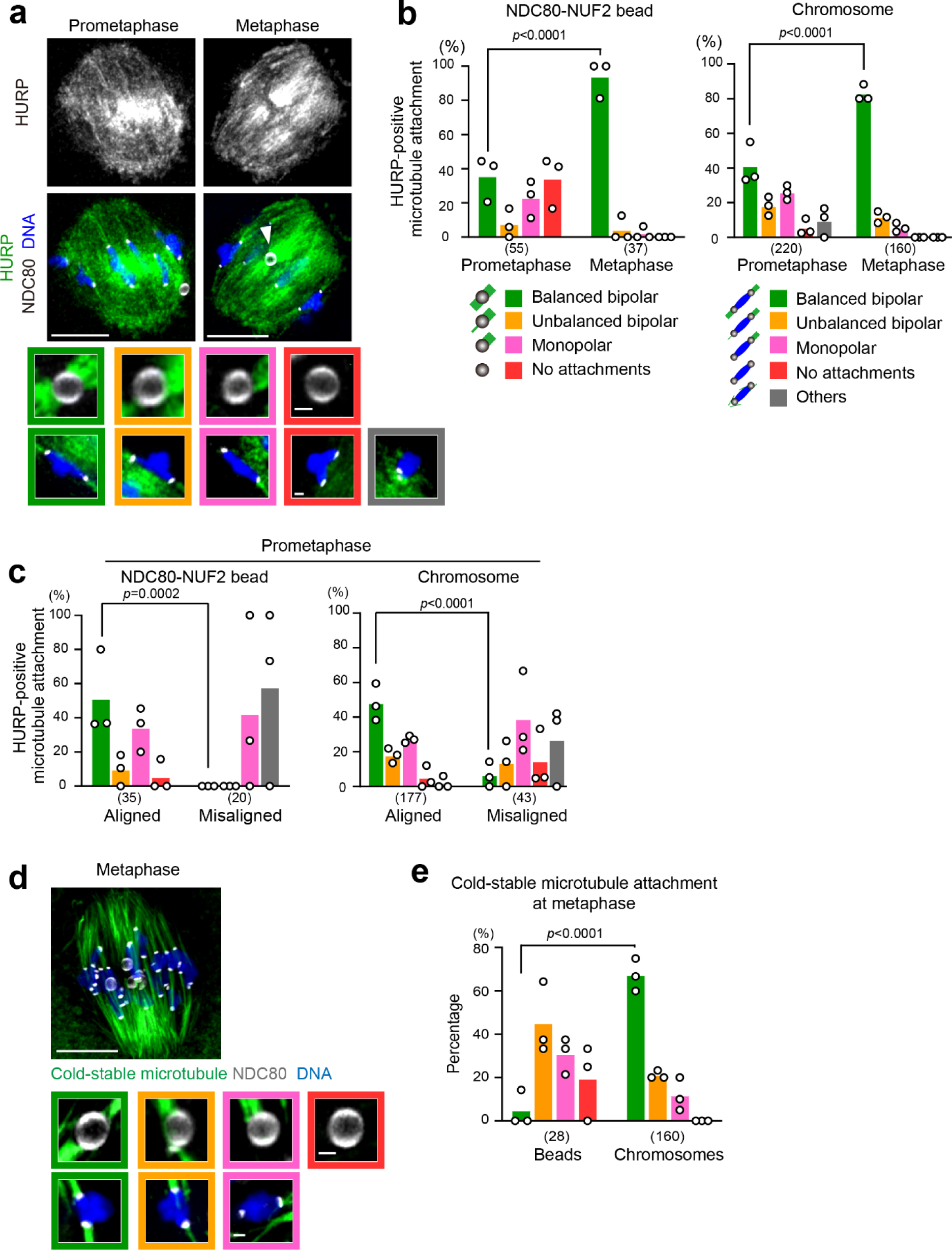
Bipolar attachment of HURP-positive microtubules. **(a)** Bipolar attachment on aligned microbeads. Confocal single-slice images of oocytes carrying NDC80-NUF2 beads, immunostained for NDC80-GFP (kinetochores and NDC80-NUF2 beads, white), HURP (green), and DNA (Hoechst33342, blue) at 3 hours (prometaphase) and 5 hours (metaphase) after NEBD. The white arrowhead indicates an NDC80-NUF2 bead with bipolar attachment. Note that the signals of NDC80-GFP uniformly located on the bead surface appear as a pair of crescents with confocal laser scanning microscopy. Scale bar, 10 µm. Attachment of HURP bundles with beads and chromosomes are categorized and magnified: balanced bipolar attachments (green frame), unbalanced bipolar attachments (orange frame), monopolar attachments (magenta frame), no attachments (red frame), and others (black frame). Scale bars, 1 µm. **(b, c)** Quantification of attachments. The number of chromosomes and beads are indicated in parentheses (from 11 prometaphase and 8 metaphase oocytes, respectively, in 3 independent experiments). Chromosomes and beads within 5 µm from the spindle equator were categorized as “Aligned”, and the others as “Misaligned”. Each dot shows the data of one experiment. P-values were calculated with Fisher’s exact test. **(d)** No prominent cold-stable microtubules on aligned microbeads. Z-projection image of metaphase oocytes carrying NDC80-NUF2 beads. Oocytes were cold-treated for 5 minutes. Cold-stable microtubules (green), NDC80-GFP (kinetochores and NDC80-NUF2 beads, white), and DNA (Hoechst33342, blue) are shown as in **(a)**. **(e)** Quantification of cold-stable attachments. Microtubule attachments are categorized and analyzed as in **(b).** The number of chromosomes and beads is indicated in parentheses (from 8 oocytes in 3 independent experiments).

### Biorientation-like state with no detectable cold-stable microtubule fibers

Kinetochore fibers, microtubule bundles with end-on attachment to kinetochores, are often visualized as a microtubule population that persists after cold treatment. In oocytes, cold-stable kinetochore fibers are rarely observed at early metaphase, when most chromosomes are aligned, and then gradually increase over time during metaphase ^25,26^. Whether the increase in cold-stable kinetochore fibers is required for the maintenance of stable alignment during metaphase is unknown. We found that at metaphase, when the vast majority of chromosomes and NDC80-NUF2 microbeads were aligned, only 7% (2/28) of NDC80-NUF2 microbeads were attached to cold-stable microtubule fibers, whereas 68% (109/160) of chromosomes were (Fig. 2d, e). These results suggest that NDC80-NUF2 microbeads are defective in forming attachment with cold-stable microtubule bundles, which is dispensable for maintaining a biorientation-like state.

### Alignment independent of SPC24-SPC25, KNL1, the MIS12 complex, and inner kinetochore proteins

Chromosome biorientation requires many outer and inner kinetochore proteins and inner centromere proteins (Fig. 3a). We therefore speculated that NDC80-NUF2 microbeads acquire a biorientation-like state by recruiting kinetochore and centromere proteins from the endogenous pool. NDC80-NUF2 microbeads recruited SPC24 and SPC25, outer kinetochore proteins that bind directly to NDC80-NUF2 (Fig. 3b and Fig. S3a), but no detectable levels of the outer kinetochore proteins DSN1, NNF1 (components of the MIS12 complex), or KNL1 (Fig. S3b). PRC1, which is recruited to the outer kinetochore via NDC80 in mouse oocytes, was found on NDC80-NUF2 microbeads (Fig. S3b), consistent with our previous report ^27^.

**Fig. 3:**
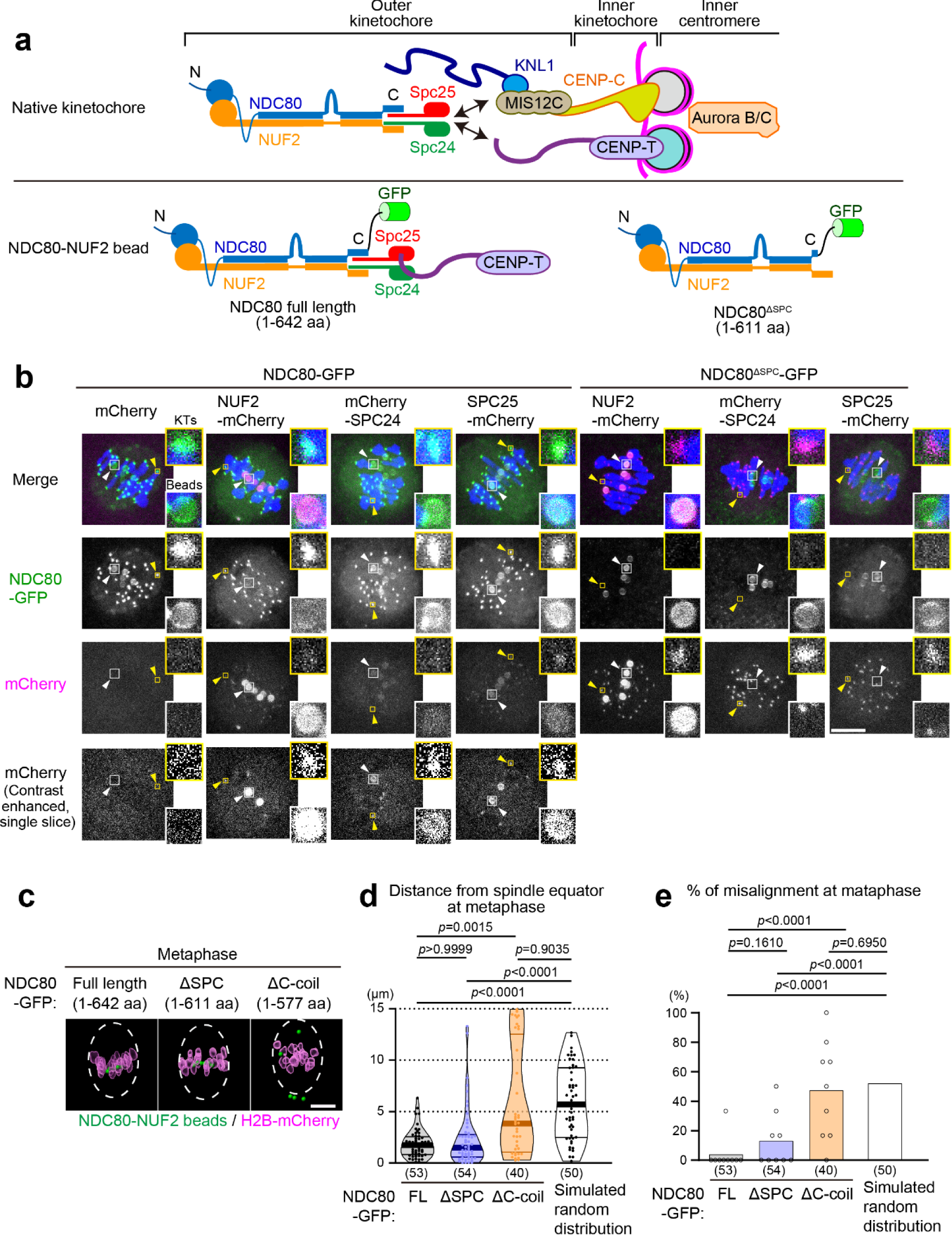
Alignment independent of SPC24-SPC25, KNL1, MIS12 complex, the inner kinetochore, and Aurora. **(a)** Composition of native kinetochores and NDC80-NUF2 beads. In the native kinetochore, NDC80-NUF2 binds to SPC24-SPC25, which binds to the MIS12 complex (MIS12C) and CENP-T. MIS12C binds to KNL1 and CENP-C. The NDC80-NUF2-SPC24-SPC25 complex, KNL1, and MIS12C forms the outer kinetochore KMN network ^10^. The inner kinetochore proteins CENP-T and CENP-C link the outer kinetochore to centromeric nucleosomes. Aurora B and its germ-cell-specific homologue Aurora C ^30^ are enriched at the inner centromere. NDC80-NUF2 beads recruit SPC24-SPC25 and CENP-T from the endogenous pool, whereas NDC80^ΔSPC^-NUF2 beads do not, as shown in **(b)** and **Fig. S3**. **(b)** NDC80-NUF2 beads recruit SPC24-SPC25, but NDC80^ΔSPC^-NUF2 beads do not. Indicated proteins were expressed in oocytes carrying anti-GFP beads. Untagged NUF2 was co-expressed unless NUF2-mCherry was used. Oocytes were fixed 5 hours after NEBD and immunostained for NDC80-GFP (green), mCherry (magenta) and DNA (Hoechst33342, blue). Z-projection images are shown unless indicated as ‘single slice’. Yellow and white arrowheads indicate kinetochores (KTs) and beads, respectively. KTs and beads are magnified. Scale bar, 10 µm. For quantification, see **Fig. S3a**. **(c)** NDC80^ΔSPC^-NUF2 beads have alignment capability. 3D-reconstructed images of chromosomes (H2B-mCherry, magenta) and beads (NDC80-GFP, green) at 6 hours after NEBD. In contrast to NDC80-NUF2^ΔSPC^ beads, NDC80-NUF2^ΔC-coil^ beads failed to align. Dashed lines indicate estimated spindle shape (20 µm in width and 30 µm in length). Scale bar, 10 µm. **(d)** Alignment of NDC80^ΔSPC^-NUF2 beads. Distance of beads from the spindle equator was quantified. The number of beads (from 9, 9, and 9 oocytes, respectively, in 3 independent experiments) is indicated in parentheses. Simulated random distribution of particles within a spindle-like ellipsoid (30 µm in length and 20 µm in width) is used as a reference. P-values were calculated with Kruskal-Wallis test with Dunn’s correction. **(e)** Misalignment frequency. “Misaligned” refers to positions over 5 µm from the spindle equator. Each dot shows the fraction in one oocyte. P-values were calculated with Fisher’s exact test.

SKA1 and SKA3, the components of the outer kinetochore SKA complex that directly interacts with NDC80-NUF2 in somatic cells ^22,28^, were enriched on microbeads but hardly detected on endogenous kinetochores in oocytes (Fig. S3c). chTOG, which interacts with NDC80-NUF2 in somatic cells ^16,17^, was not detected on endogenous kinetochores in oocytes, as reported previously ^29^, nor on NDC80-NUF2 microbeads (Fig. S3d). The inner kinetochore protein CENP-T, but neither CENP-C nor the inner centromere protein Aurora B nor C ^30^, was detected on NDC80-NUF2 microbeads (Fig. S3e–g). We conclude that NDC80-NUF2 microbeads establish stable alignment independently of the recruitment of KNL1, the MIS12 complex, CENP-C, and Aurora B and C kinases (Fig. 3a).

SPC24 and SPC25 form a heterodimer that bridges the C-terminal domains of NDC80-NUF2 to the MIS12 complex or the inner kinetochore CENP-T (Fig. 3a) ^1,2^. To investigate whether SPC24-SPC25 and its interactors are required for the biorientation-like state of NDC80-NUF2 microbeads, we disrupted the interaction between NDC80-NUF2 and SPC24-SPC25 by deleting a short segment of the C-terminus (a.a. 612–642) of NDC80 (the resulting form is hereafter called NDC80^ΔSPC^) (Fig. 3a). NDC80^ΔSPC^-NUF2 microbeads did not contain any detectable SPC24-SPC25 or CENP-T (Fig. 3b, Fig. S3a, e). Consistently, NDC80^ΔSPC^-GFP did not localize to endogenous kinetochores (Fig. S4). Nevertheless, NDC80^ΔSPC^-NUF2 microbeads established alignment, similar to NDC80-NUF2 microbeads (Fig. 3c–e, see below), whereas NDC80^ΔC-coil^-NUF2, in which a larger segment of the C-terminus (a.a. 578–642) of NDC80 was deleted, failed to align microbeads (Fig. 3c–e). These results suggest that NDC80-NUF2 microbeads establish a biorientation-like state independently of the recruitment of SPC24-SPC25 and its inner kinetochore interactors such as CENP-T.

### Alignment independent of NDC80 phosphoregulation

Chromosome biorientation depends on Aurora kinase-mediated phosphorylation of the N-terminal tail of NDC80, which decreases the microtubule-binding affinity of NDC80-NUF2 for attachment error correction ^3,4,8,10,11^. To ask whether the phosphoregulation of the NDC80 tail is required for NDC80-NUF2 microbead alignment, we used the phospho-mutant forms of NDC80^ΔSPC^, which does not localize to, and thus does not affect, endogenous kinetochores. Consistent with the idea that NDC80-NUF2 microbeads establish alignment through their microtubule-binding activity, NDC80^ΔSPC^-9D-NUF2-coated microbeads, which carry phospho-mimetic mutations, failed to establish alignment (Fig. S5a). In contrast, NDC80^ΔSPC^-9A-NUF2 microbeads, which carry phospho-deficient mutations, established alignment with comparable kinetics to NDC80^ΔSPC^-NUF2 microbeads (Fig. 4a–c, Supplementary Video 5).

**Fig. 4:**
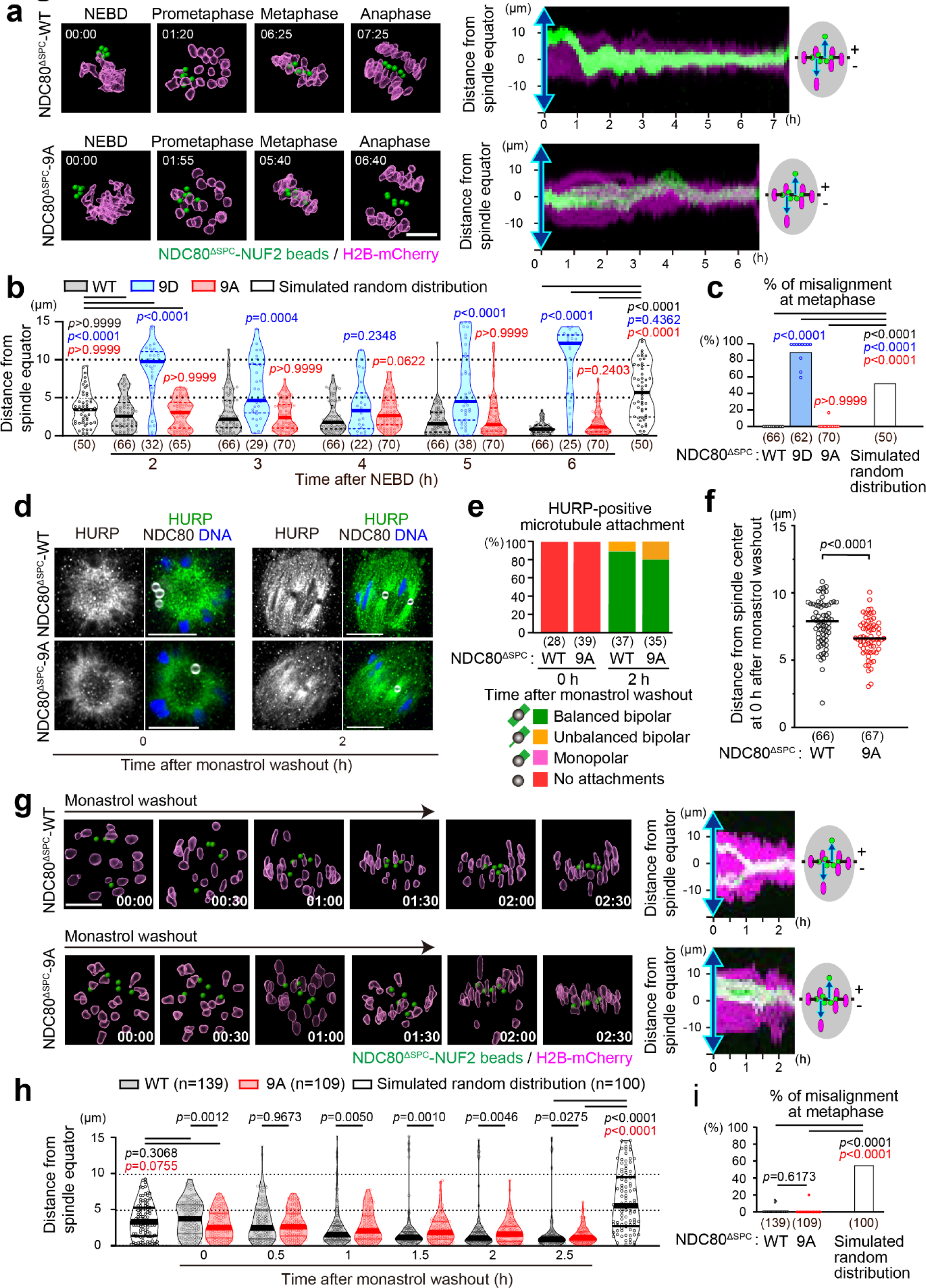
Alignment independent of NDC80 phosphoregulation. **(a)** Live imaging. 3D-reconstructed images of chromosomes (H2B-mCherry, magenta) and NDC80^ΔSPC^-WT/9A-NUF2 beads (green). Time after NEBD (hh:mm). Scale bar, 10 µm. The kymograph shows projected signals on the spindle axis over time. The vertical axis shows distance from the spindle equator. Images for NDC80^ΔSPC^-9D-NUF2 acquired in the same experiment are shown in **Fig. S5a**. **(b)** Alignment efficiency. Distance of NDC80^ΔSPC^-WT/9A/9D-NUF2 beads from the spindle equator was quantified. The number of beads (from 11, 11, and 12 oocytes, respectively, in 3 independent experiments) is indicated in parentheses. Simulated random distribution of particles within a spindle-like ellipsoid (20 µm at 2h or 30 µm at 6 h in length and 20 µm in width) is used as a reference. NDC80^ΔSPC^-9D-NUF2 beads apparently positioned outside the spindle (>10 µm from the spindle axis or >15 µm from the equator) were excluded from the analysis. P-values against WT or simulated values were calculated using Kruskal-Wallis test with Dunn’s correction. **(c)** Misalignment frequency. “Misaligned” refers to positions over 5 µm from the spindle equator at 6 hours after NEBD. Each dot shows the data of an oocyte. P-values were calculated with Fisher’s exact test. **(d)** Microtubule attachment. Confocal single-slice images of oocytes after monastrol washout, immunostained for GFP (NDC80^ΔSPC^-GFP, white), HURP (green), and DNA (Hoechst33342, blue). Scale bar, 10 µm. **(e)** Attachments with microtubule bundles in **(d)** are categorized and quantified. The number of beads (from 7, 9, 9, and 9 oocytes, respectively, in 3 independent experiments) is shown in parentheses. **(f)** NDC80^ΔSPC^-9A-NUF2 beads are located inwardly. In monastrol-treated oocytes, the positions of NDC80^ΔSPC^-WT/9A-NUF2 beads were analyzed. The number of beads (from 26, 20, oocytes, respectively, in 3 independent experiments) is shown in parentheses. P-value was calculated using unpaired Welch’s t-test. **(g)** Live imaging. 3D-reconstructed images of chromosomes (H2B-mCherry, magenta), NDC80^ΔSPC^-WT/9A-NUF2 beads (green) after monastrol washout. Time after monastrol washout (h:mm). Scale bar, 10 µm. Kymographs along the spindle axis are shown. **(h)** Alignment efficiency. Distance of NDC80^ΔSPC^-WT/9A-NUF2 beads from the spindle equator after monastrol washout was quantified as in **(b)**. The number of beads (from 28 and 22 oocytes, respectively, in 3 independent experiments) is indicated in parentheses. P-values were calculated using Mann-Whitney test. **(i)** Misalignment frequency. “Misaligned” refers to positions over 5 µm from the spindle equator at 2.5 hours after monastrol washout. Each dot shows the data of an oocyte. P-values were calculated with Fisher’s exact test.

Consistently, NDC80-NUF2 microbeads established alignment in oocytes treated with AZD1152, an inhibitor of both Aurora B and C ^26^, while chromosomes exhibited misalignment (Fig. S5b–d). These results demonstrate that Aurora kinase-mediated phosphorylation of the NDC80 tail is dispensable for the alignment of NDC80-NUF2 microbeads.

### NDC80 phosphoregulation promotes a biorientation-like state under a challenging condition

A most stringent test for the capacity of biorientation is a monastrol washout assay ^31^. In this assay, cells are treated with monastrol, a drug that induces monopolar spindle formation, thereby increasing the likelihood of erroneous kinetochore-microtubule attachments. Monastrol is then washed out and the ability of biorientation is assessed.

In monastrol-treated oocytes, NDC80^ΔSPC^-NUF2 microbeads were located around the surface of the monopolar spindle, similar to chromosomes, with no attachment to HURP-positive microtubule bundles (Fig. 4d, e, “0 h”). Notably, NDC80^ΔSPC^-9A-NUF2 microbeads were significantly more inwardy located (Fig. 4f), although attachment to HURP-positive microtubule bundles was not observed (Fig. 4d, e, “0h”). These results suggest that NDC80-NUF2 microbeads suppress monopolar attachment of HURP-negative microtubules via NDC80 tail phosphorylation.

Upon monastrol washout, NDC80^ΔSPC^-NUF2 microbeads gradually aligned, similar to chromosomes (Fig. 4g–i, Supplementary Video 5). The aligned microbeads were attached with bipolar HURP-positive microtubule bundles (Fig. 4d, e, “2 h”). Remarkably, NDC80^ΔSPC^-9A-NUF2 microbeads also established alignment by 2.5 hours after monastrol washout (Fig. 4g–i), resulting in the formation of bipolar attachment with HURP-positive microtubule bundles (Fig. 4d, e). Thus, NDC80 tail phosphorylation is dispensable for establishing a biorientation-like state. However, quantitative analysis showed that the alignment of NDC80^ΔSPC^-9A-NUF2 microbeads was significantly delayed compared to NDC80^ΔSPC^-NUF2 microbeads (Fig. 4h). Thus, the phosphoregulation of the NDC80 tail is required for NDC80-NUF2 microbeads to efficiently establish a biorientation-like state for recovery from challenging conditions.

### Large platform size promotes alignment

The size of the NDC80-NUF2 microbeads used in the above experiments was 1.7–2.6 µm in diameter, which is one order of magnitude larger than endogenous kinetochores (∼250 nm wide) ^32,33^. We reasoned that the large platform of the microbeads for NDC80-NUF2 recruitment promoted their establishment of a biorientation-like state. To test this possibility, we co-injected microbeads of smaller sizes, ranging from 0.4 to 1.7 µm in diameter, together with larger 1.7–2.6 µm microbeads into oocytes, and compared their alignment efficiencies within the same oocytes by live imaging (Fig. 5a). Notably, this analysis revealed that smaller NDC80-NUF2 microbeads exhibited a significant delay in alignment compared to larger microbeads (Fig. 5b–d). These results suggest that the NDC80-NUF2-mediated biorientation mechanism is enhanced by increased size of its platform.

**Fig. 5:**
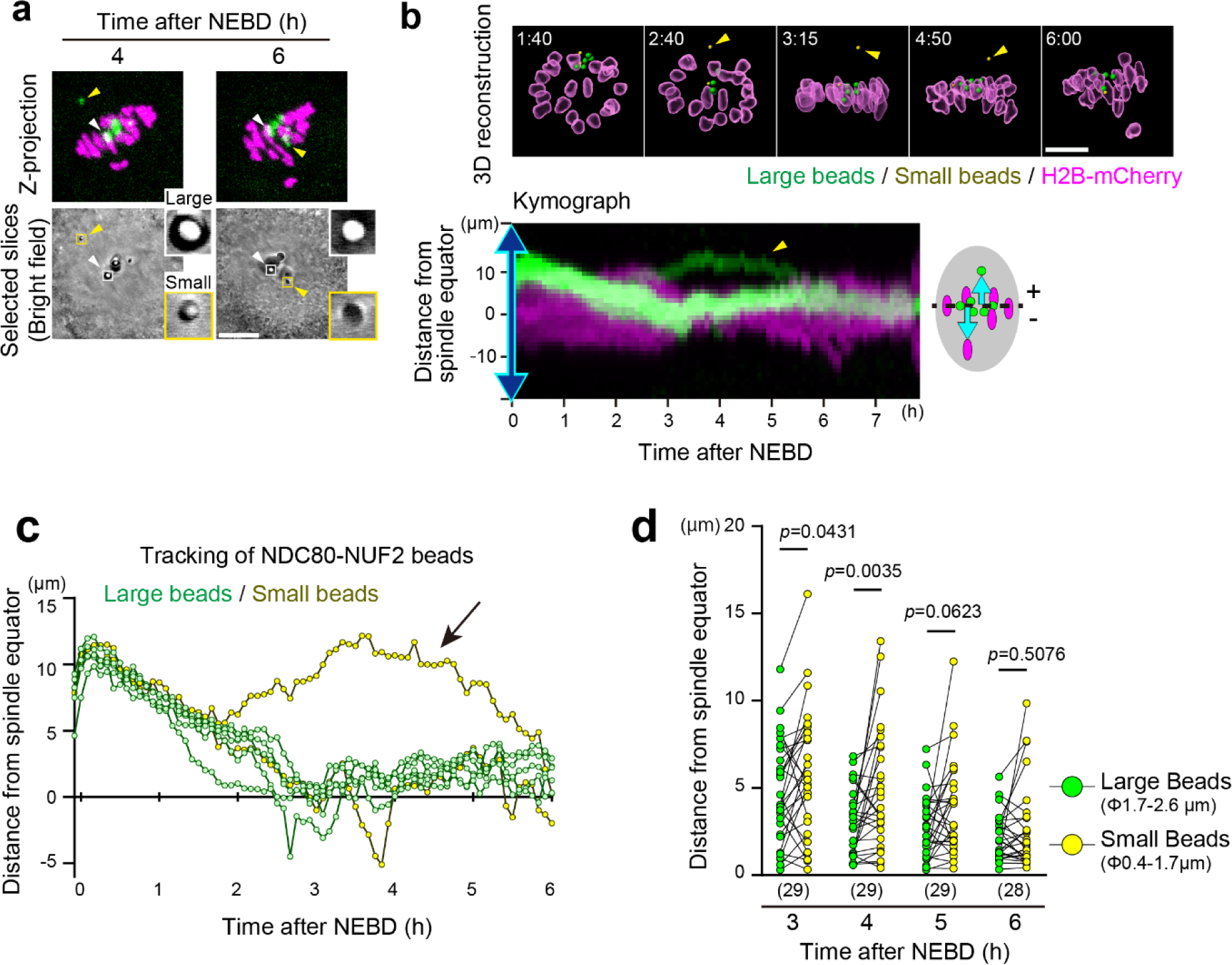
Large platform size promotes NDC80-NUF2 microbead alignment. **(a)** NDC80-NUF2 beads of different sizes. NDC80-GFP and NUF2 were expressed in oocytes carrying anti-GFP beads of different sizes. Z-projection and selected slice images of NDC80-GFP (green) and H2B-mCherry (magenta) are shown. White and yellow arrowheads indicate beads of 1.9 µm (“Large”) and 1.1 µm (“Small”) in diameter, respectively, which are magnified in insets. Scale bar, 10 µm. **(b)** Smaller NDC80-NUF2 microbeads exhibit less efficient alignment. 3D-reconstructed images of chromosomes (H2B-mCherry, magenta) and NDC80-NUF2 beads of different sizes (large beads, green; small beads, yellow, indicated by arrowheads). Time after NEBD (h:mm). Scale bar, 10 µm. Kymographs were generated with projected signals based on the spindle axis. The vertical axis shows the distance from the spindle equator. Arrows indicate a small bead (1.1 µm in diameter). **(c)** Tracking. Images in **(b)** were used for tracking. **(d)** Alignment efficiency. The distances of NDC80-NUF2 beads from the spindle equator were measured. Each dot indicates the average value of beads within one oocyte. Lines connect the data from the same oocytes. The number of oocytes from 3 independent experiments is indicated in parentheses. P-values were calculated with Wilcoxon matched-pairs signed rank test.

## Discussion

Here we have demonstrated that an artificial kinetochore-like microbead, anchoring the outer kinetochore NDC80-NUF2 heterodimer, is capable of establishing a biorientation-like state within the spindle of live mouse oocytes. Previous studies in budding yeast have shown that tethering the fungal outer kinetochore Dam1/DASH complex to sister DNAs allows biorientation depending on other kinetochore proteins and Aurora ^34,35^. Studies using vertebrate cells induced ectopic kinetochore assembly with the inner kinetochore protein CENP-T tethered to sister DNAs, which partially supported chromosome segregation ^36^. Most recently, a study shows that genetically encoded nanoscale particles tethering CENP-T recruits kinetochore proteins capable of interacting with microtubules ^37^. Our study used a biohybrid approach with a microscale bead tethering NDC80-NUF2 on its surface, showing the emergence of the capacity to establish a biorientation-like state by bypassing the requirement for the outer kinetochore proteins SPC24-SPC25, KNL1 and the MIS12 complex, the inner kinetochore proteins CENP-T and CENP-C, the inner centromere kinase Aurora, and DNA.

These results suggest that the NDC80-NUF2 branch of the outer kinetochore has an intrinsic mechanism sufficient to establish a biorientation-like state. This property depends on the middle loop and C-terminal coiled-coil domains of NDC80 (Fig. 3c–e, Fig. S1b), which mediate protein-protein interaction ^21–23,27,28^. In somatic cells, chTOG interaction with the NDC80 complex mediates tension-dependent stabilization of kinetochore-microtubule attachment ^16,17^. However, chTOG enrichment was not observed on NDC80-NUF2 microbeads (Fig. S3d). Thus, interaction with other proteins or the intrinsic property of NDC80-NUF2 likely drives a catch bond-like behavior to directly stabilize microtubule attachment upon tension ^15,16,38^, establishing a biorientation-like state. Our results also suggest that NDC80 phosphorylation, likely by cytoplasmic Aurora, contributes to efficient biorientation after monastrol washout. The NDC80-NUF2 branch may intrinsically regulate Aurora activity or its effect in a tension-dependent manner ^39^.

Our observations shed light on physical requirements for biorientation. First, microbeads without a bipolar cue can robustly achieve a biorientation-like state, indicating that the presence of two physically separate platforms and their geometry for microtubule attachment, like the sister kinetochores of the chromosome, is not a prerequisite for establishing bipolar attachment. This view is consistent with the phenomenon of merotelic attachments, where one kinetochore is attached by microtubules from opposite spindle poles, often leading to aneuploidy, particularly in cancer cells ^40^. Second, alignment efficiency depends on microbead size, implying that the NDC80-NUF2-mediated biorientation benefits from an expanded platform for outer kinetochore assembly. Such platform expansion may be attributed to the evolution of the inner kinetochore, which possesses self-oligomerization activity ^41,42^, and the centromere, which contains active DNA arrays that extend ∼0.3–4.8 Mbps in humans ^43^. The platform of microbeads that we used in this study is an order of magnitude larger than the native kinetochore, which we suggest helps the establishment of a biorientation-like state by the NDC80-NUF2 branch alone.

Although NDC80-NUF2 microbeads functionally resemble chromosomes in terms of efficient and robust alignment and bipolar microtubule attachment, they are distinct in the capacity to mature microtubule attachment into a cold-stable state (Fig. 2d, e). Maturation into cold-stable microtubule attachment may require additional factors such as the outer kinetochore proteins KNL1 and the MIS12 complex ^10^, inner kinetochore proteins, or mechanisms such as spatial regulation of NDC80 phosphorylation ^3,4,8^. Moreover, NDC80-NUF2 microbeads are preferentially located in the inner region of the metaphase plate (Fig. 1b, c, Fig. S1d), perhaps due to the lack of chromokinesin-mediated polar ejection force ^44^. Future studies aimed at artificially constructing kinetochores and chromosomes will provide a further understanding of the respective roles of kinetochore, centromere and chromosome proteins to the biorientation process. Kinetochore-like microbeads, together with chromosome-like microbeads capable of inducing bipolar spindle self-assembly ^45^, hold promise as tools for advancing the development of synthetic DNA segregation.

In summary, our biohybrid kinetochore approach has identified a minimal kinetochore branch that enables an apolar microscale particle to achieve biorientation in live cells. This finding may serve as a foundation for the development of microscale machines or artificial organelles designed to autonomously segregate materials in living systems.

## Methods

### Mice

All animal experiments were approved by the Institutional Animal Care and Use Committee at RIKEN Kobe Branch (IACUC). Mice were housed in 12-hour light/12-hour dark cycle in temperature of 18–23 °C with 40–60% humidity environment. B6D2F1 (C57BL/6 × DBA/2) female mice, 8–12 weeks of age, were used to obtain oocytes.

### Mouse oocyte culture

Mice were intraperitoneally injected with 0.1 ml of CARD HyperOva (KYUDO) or 0.2 ml (10 IU) of pregnant mare serum gonadotrophin (PMSG). Fully-grown oocytes at the germinal vesicle (GV) stage were collected 48 hours after injection and released into M2 medium containing 200 µM 3-isobutyl-1-methylxanthine (IBMX, Sigma) at 37 °C. Meiotic resumption was induced by washing to remove IBMX. When indicated, 100 µM monastrol (Sigma), 5 µM ProTAME (Bio-Techne R&D Systems) and 500 nM AZD1152 (Cosmo-bio) were used. Monastrol was added 30 minutes after IBMX and washed out 2 hours after NEBD.

### RNA microinjection

mRNAs were transcribed *in vitro* using the mMESSAGE mMACHINE T7 kit (Thermo Fisher Scientific) and purified. The synthesized RNAs were stored at −80°C until use. The mRNAs were introduced into fully-grown GV-stage mouse oocytes through microinjection. The microinjected oocytes were cultured at 37 °C for 2 hours before microbead microinjection. Microinjections were performed with 1 pg of NDC80(WT/9A/9D/ΔSPC/ΔC-Coil/ΔLoop/DFAA)-GFP, 1 pg of NUF2, 1 pg of NUF2-mCherry, 1.5 pg of H2B-mCherry, 3.2 pg of KNL1-GFP, 1 pg of MIS12-GFP, and 1 pg of GFP, 1 pg of mCherry-SPC24, 1 pg of SPC25-mCherry, 1 pg of CENPT-mCherry, 1.2 pg of mCherry-SKA1, 1.2 pg of mCherry-SKA3, 1.2 pg of AURKB-mCherry, 1.2 pg of AURKC-mCherry and 1.2 pg of mCherry.

### Bead microinjection

Anti-GFP mAb-Magnetic Beads (2.0 μm in diameter, D153-11, MBL) were introduced into mRNA injected GV-stage oocytes through microinjection with a piezo pulse (Supplementary Video 1). For most experiments, 5–7 beads of 1.7–2.6 μm in diameter were microinjected into each oocyte. When indicated, we selectively picked up a total of 5–7 beads, including 1-4 smaller beads (0.4–1.7 μm in diameter) and 2–5 larger beads (1.7-2.6 μm in diameter) and microinjected them into each oocyte (Fig. 5). The microinjected oocytes were cultured at 37 °C for 1 hour before meiotic resumption.

### Live imaging

An LSM880 confocal microscope (Carl Zeiss) equipped with a 40×C-Apochromat 1.2NA water immersion objective lens (Carl Zeiss) was controlled using Zen software with the multiposition autofocus macro PipelineConstructor ^46^. For imaging of chromosomes and beads, we recorded 21–25 z-confocal sections (every 1.25–1.5 μm) of 512 × 512 pixel or 256 × 256 pixel xy images, which covered a total volume of at least 32.7 μm × 32.7 μm × 30 μm, at 5-minute time intervals for at least 12 hours after the induction of maturation.

### Immunostaining

Oocytes were fixed in fixation buffer (100 mM PIPES (pH 7.0), 1 mM MgCl_2_, 0.1% Triton-X100) plus 1.6% formaldehyde for 30 minutes at room temperature, washed four times with PBT, and blocked with 3% BSA in PBT for incubated overnight at 4℃. Oocytes were incubated with primary antibodies in 3% BSA-PBT overnight at 4°C, washed with 3% BSA-PBT, and then incubated with secondary antibodies and 5 μg/ml Hoechst33342 for 2 hours at room temperature or overnight at 4 ℃. Primary antibodies used were rabbit anti-KNL1 (ab70537, abcam; 1:50), rabbit anti-DSN1 (gift from Dr. H. Shibuya; 1:500), rabbit anti-NNF1(gift from Dr. H. Shibuya; 1:500), rabbit anti-PRC1 (sc-8356, Santa Cruz Biotechnology; 1:100), rabbit anti-CENP-C (gift from Dr. Y. Watanabe; 1:500), rabbit control IgG (12-370, sigma-aldrich; 1:100), mouse anti-chTOG (cosmo bio, 67631-1-IG; 1:500), rabbit anti-GFP (ab6556, abcam; 1:500), mouse anti-mCherry (632543, Clontech; 1:500), mouse anti-GFP (ab1218, abcam; 1:500), rabbit anti-HURP (sc-98809, Santa Cruz Biotechnology; 1:500), and rat anti-α tubulin (YL1/2) (MCA77G, Bio-Rad; 1:1500). Secondary antibodies used were Alexa Fluor 488 goat anti-mouse IgG (H + L) (A11029), goat anti-rabbit IgG (H + L) (A11034), Alexa Fluor 555 goat anti-rat lgG (H+L) (A21434), Alexa Fluor 568 goat anti-mouse IgG (H + L) (A11031), goat anti-rabbit IgG (H + L) (A11036), Alexa Fluor 647 goat anti-Human IgG (H + L) (A21445), and donkey anti-rabbit lgG (H+L) (A31573) (1:500, Molecular Probes). The oocytes were washed and stored in 0.01% BSA-PBS. The oocytes were imaged under an LSM780 confocal microscope (Carl Zeiss) equipped with a 40×C-Apochromat 1.2NA water immersion objective lens (Carl Zeiss). We recorded at least 14 z-confocal sections (every 1.0 µm) of 512 × 512 pixel xy images, which covered a total volume of at least 26.57 μm × 26.57 μm × 13 μm. For HURP and microtubules, oocytes were imaged under an LSM880 with Airyscan confocal microscope (Carl Zeiss) equipped with a 40×C-Apochromat 1.2NA water immersion objective lens (Carl Zeiss). We recorded at least 90 z-confocal sections (every 0.2 µm) of 1024 × 1024 pixel xy images, which covered a total volume of at least 31.17 µm × 31.17 µm × 17.8 µm. For visualization of cold-stable microtubules, oocytes were incubated in an ice-cold M2 medium for 5 minutes before fixation. Note that the signals of NDC80-GFP uniformly located on the bead surface appear as a pair of crescents with confocal laser scanning microscopy.

### 4D analysis

Chromosome surface in 3D was reconstructed using Imaris software (Bitplane). Anti-GFP bead position was manually defined based on bright-field and GFP images. For every time point, the spindle axis was manually estimated based on chromosome distribution and shape. For time points from 0 to 3 hours after NEBD (before the establishment of the spindle axis), the same spindle axis as that of 4 h after NEBD was used. The spindle center was defined as the center of mass of all chromosome positions. To generate kymographs of NDC80-NUF2 beads and chromosomes, the intensities of their signals were projected onto the spindle axis or the spindle equator for each time point with an in-house macro in Fiji.

### Signal intensity quantification

Quantification of fluorescence signals on beads and kinetochores was performed in Fiji or Imaris. First, the mean fluorescence intensity on a bead or a kinetochore was measured and subtracted by the mean intensity in an adjacent cytoplasmic region. Second, the value of the protein of interest (e.g., mCherry-tagged or antibody-labeled proteins) relative to that of the reference (e.g., NDC80-GFP) was calculated.

### Statistical analysis

Graphs were generated and statistical analyses were performed using Excel and GraphPad Prism. No statistical analysis was performed to predetermine the sample size. Methods for significance tests are described in figure legends. Random distribution of particles within a spindle-like ellipsoid was simulated by an in-house macro in Fiji.

## Acknowledgements

We thank Drs. S. Yoshida and H. Kyogoku for technical assistance for microbead injection, H. Shibuya and Y. Watanabe for providing antibodies, J. Ellenberg for a microscope automation macro, the imaging and animal facilities of RIKEN BDR for technical support, and all lab members for discussions and comments. K.A. and Y.Z. are supported by the RIKEN Junior Research Associate program. This work was supported by RIKEN intramural grants, JSPS KAKENHI Grant Number 23H04948, 21H02407, and 18H05549, and a grant from the Mitsubishi Foundation to T.S.K.

## Author contributions

K.A. and T.S.K. conceived the project. K.A. performed almost all the experiments. Y.Z. performed experiments shown in Figs. 2 and 4d–i. T.S.K. supervised the project. K.A., Y.Z., and T.S.K. wrote the manuscript.

## Competing interests

The authors declare no competing financial interests.

---

**Fig. S1:**
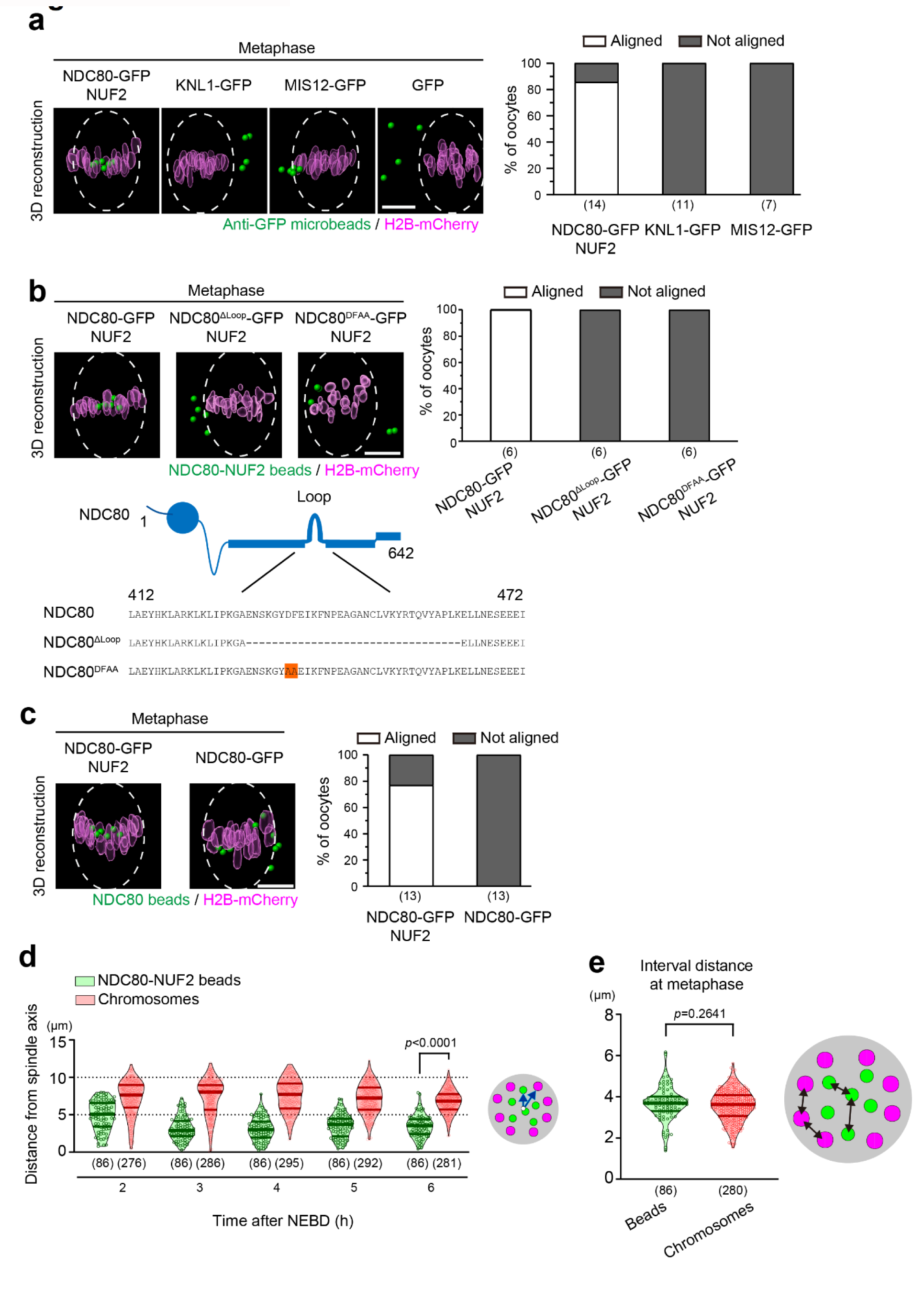
NDC80-NUF2-specific effects on alignment. **(a–c)** Experiments were designed to tether the indicated proteins to anti-GFP beads as in Fig. 1a. 3D-reconstructed images of chromosomes (H2B-mCherry, magenta) and beads (green) at metaphase (5 hours after NEBD). Dashed lines indicate estimated spindle shape (20 µm in width and 30 µm in length). Scale bar, 10 µm. Oocytes were categorized into “Aligned” when all beads were positioned within 6 µm from the spindle equator and 8 µm from the spindle axis. The number of oocytes is indicated in parentheses. In (**b**), the construct designs of NDC80 loop-deleted (NDC80^ΔLoop^) and -mutated (NDC80^DFAA^) forms ^23^ are shown at the bottom. **(d)** NDC80-NUF2 beads occupy the inner region of the metaphase plate. Distance from the spindle axis is plotted. The number of beads or chromosomes from 3 independent experiments is indicated in parentheses. P-value was calculated with Mann-Whitney’s U-test. **(e)** Neighborhood distance of NDC80-NUF2 beads is similar to that of chromosomes. Distance between neighboring beads or chromosomes at metaphase (6 h after NEBD) is plotted. The number of beads or chromosomes from 3 independent experiments is indicated in parentheses. P-value was calculated with Mann-Whitney’s U-test.

**Fig. S2:**
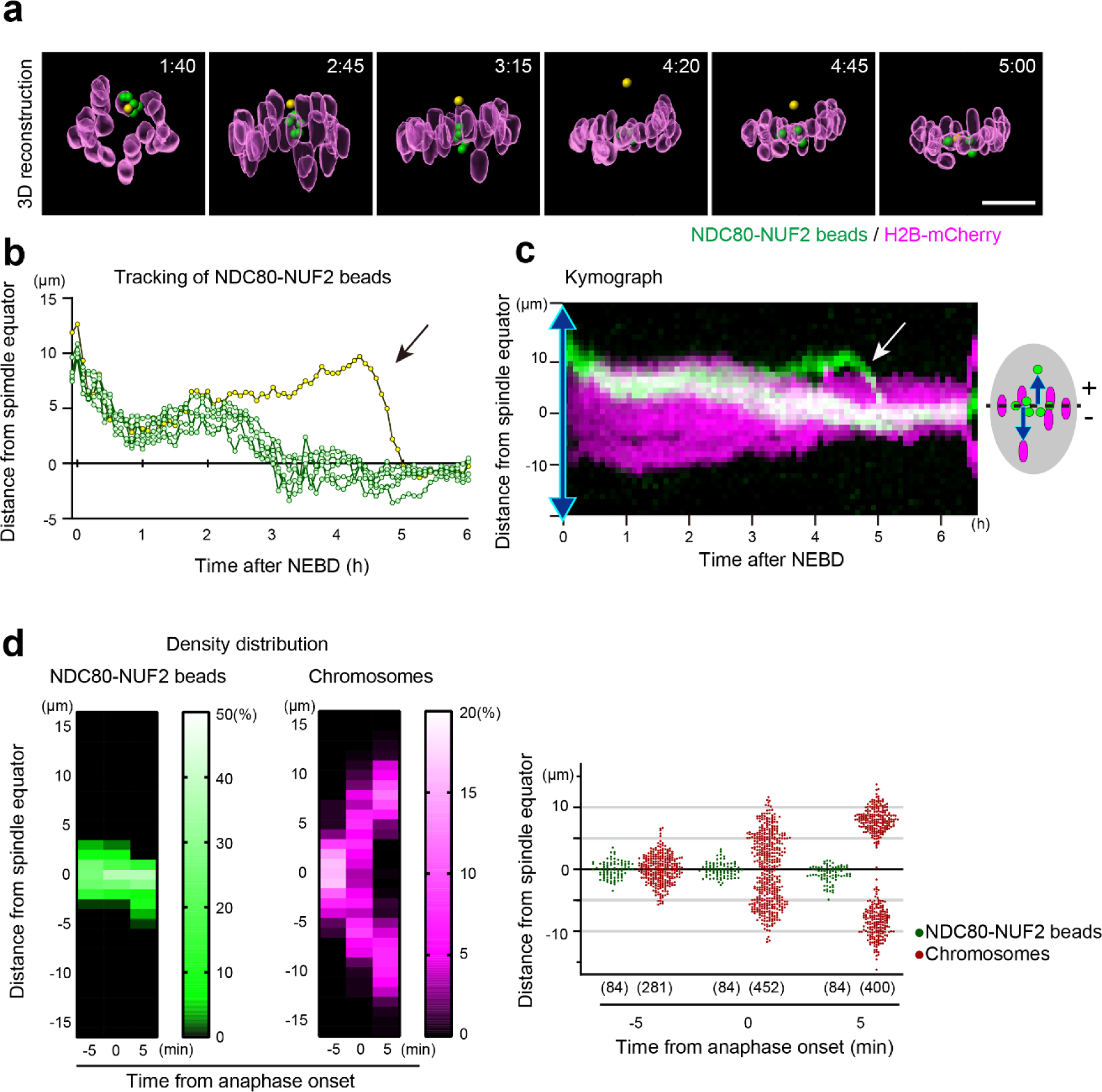
Error correction of alignment. **(a)** Live imaging. 3D-reconstructed images of chromosomes (H2B-mCherry, magenta) and NDC80-NUF2 beads (green). An NDC80-NUF2 bead exhibiting transient misalignment followed by error correction is shown (yellow). Time after NEBD (h:mm). Scale bar, 10 µm. **(b)** Tracking. NDC80-NUF2 beads in (**a**) were tracked in 3D. Distance from the spindle equator over time is shown. **(c)** Kymograph. The kymograph shows projected signals on the spindle axis over time. The vertical axis shows the distance from the spindle equator. **(d)** Alignment prior to anaphase. Average density maps (left) and positions (right) of NDC80-NUF2 beads and chromosomes, respectively, along the spindle axis. The number of beads and chromosomes from 14 oocytes in 3 independent experiments is shown in parentheses.

**Fig. S3:**
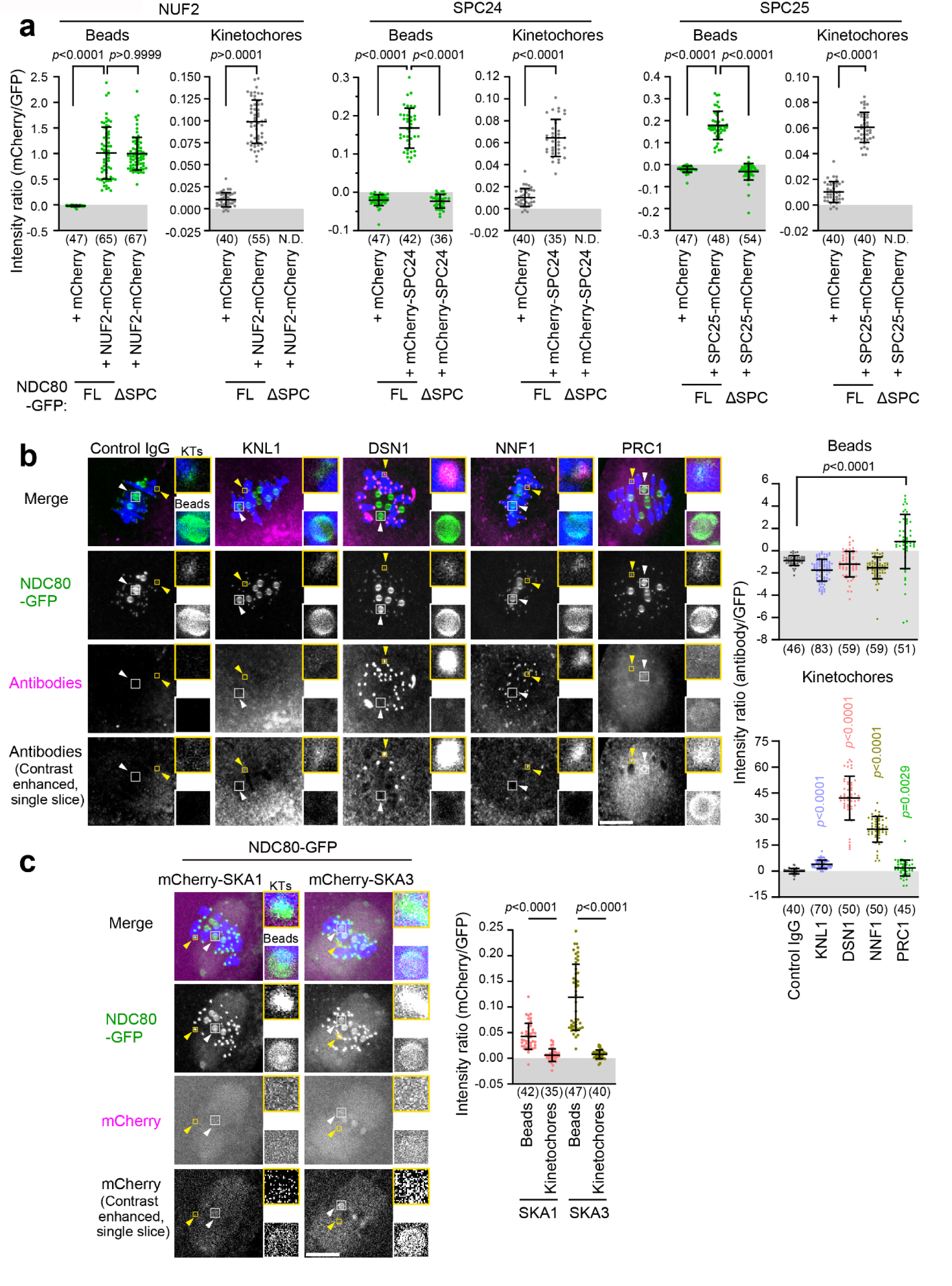
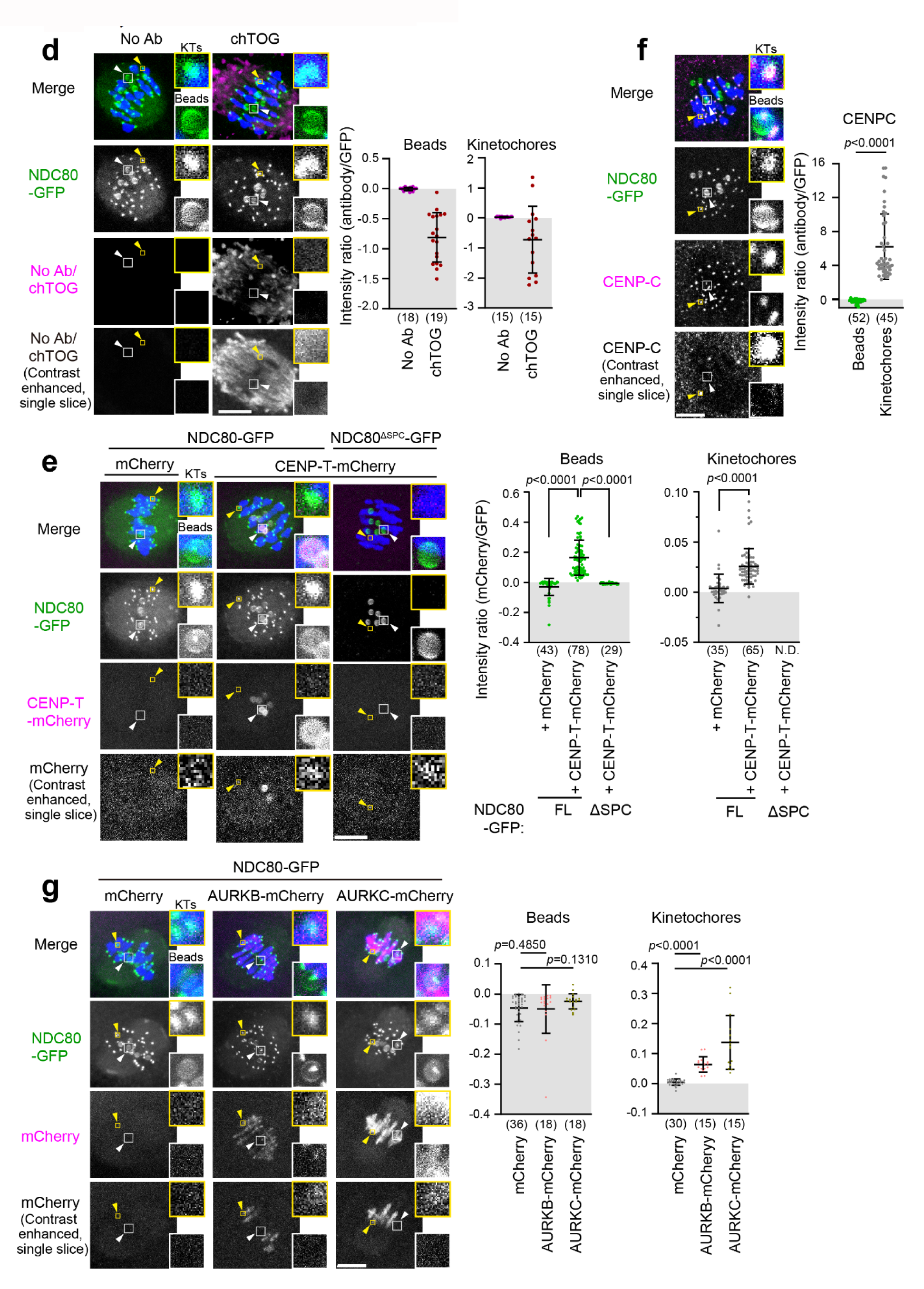
Localization of kinetochore proteins on NDC80-NUF2 beads. **(a)** Quantification of NUF2 and SPC24-SPC25 on NDC80-NUF2 beads and kinetochores. Images are shown in Fig. 3b. The cytoplasm-subtracted intensity of mCherry signals on beads or kinetochores, relative to that of NDC80-GFP signals, was calculated. The number of beads and kinetochores (each group from at least 6 oocytes in 3 independent experiments) are shown in parentheses. The datasets of ‘NDC80-FL-GFP + mCherry’ were acquired in the same experiments and are identical among the graphs. N.D., not determined due to the lack of detectable NDC80^ΔSPC^-GFP on kinetochores (**Fig. S4**). P-values were calculated with Kruskal-Wallis test with Dunn’s correction on beads and Mann-Whitney’s U-test on kinetochores. **(b)** The outer kinetochore proteins KNL1, DSN1, and NNF1 are enriched on kinetochores but not on NDC80-NUF2 beads. Oocytes were immunostained 5 hours after NEBD for NDC80-GFP (green), indicated proteins (KNL1, DSN1, NNF1, PRC1 and control IgG antibodies, magenta) and DNA (Hoechst33342, blue). Yellow and white arrowheads indicate kinetochores (KTs) and beads, respectively. Kinetochores and beads are magnified. Z-projection images are shown unless indicated as ‘single slice’. The intensity ratio was calculated as in **(a)**. The number of beads and kinetochores (each group from at least 8 oocytes in at least 3 independent experiments) are shown in parentheses. Scale bar, 10 µm. P-values were calculated with Mann-Whitney’s U-test and Kruskal-Wallis test with Dunn’s correction on beads and kinetochores, respectively. **(c)** The NDC80 interactor SKA1 and SKA3 are enriched on NDC80-NUF2 beads but hardly detected on kinetochores. Indicated proteins were expressed in oocytes carrying anti-GFP beads. Oocytes were fixed 5 hours after NEBD and immunostained for GFP (green), mCherry (magenta) and DNA (Hoechst33342, blue). Images are shown and signals are quantified as in **(b)**. The number of beads and kinetochores (each group from at least 7 oocytes in 2 independent experiments) are shown in parentheses. P-values were calculated with Mann-Whitney’s U-test. **(d)** chTOG is not enriched on NDC80-NUF2 beads or kinetochores. chTOG was analyzed and images are shown as in **(c)**. The number of beads and kinetochores (each group from 3 oocytes) is shown in parentheses. **(e)** CENP-T is enriched on NDC80-NUF2 beads but not on NDC80^ΔSPC^-NUF2 beads. CENP-T-mCherry was analyzed and images are shown as in **(b)**. The number of beads and kinetochores (from at least 5 oocytes) are shown in parentheses. P-values were calculated with Kruskal-Wallis test with Dunn’s correction and Mann-Whitney’s U-test on beads and kinetochores, respectively. **(f)** CENP-C is not enriched on NDC80-NUF2 beads. Oocytes were fixed 5 hours after NEBD and immunostained for NDC80-GFP (green), CENP-C (magenta), and mCherry (H2B-mCherry, blue). CENP-C was analyzed and images are shown as in **(b)**. The number of beads and kinetochores (from 9 oocytes in 3 independent experiments) are shown in parentheses. P-value was calculated with Mann-Whitney’s U-test. **(g)** Aurora B or C is not enriched on NDC80-NUF2 beads. AURKB-mCherry and AURKC-mCherry were analyzed and images are shown as in **(b)**. The number of beads and kinetochores (from 6, 3 and 3 oocytes, respectively, in 2 independent experiments) are shown in parentheses. P-values were calculated with Kruskal-Wallis test with Dunn’s correction.

**Fig. S4:**
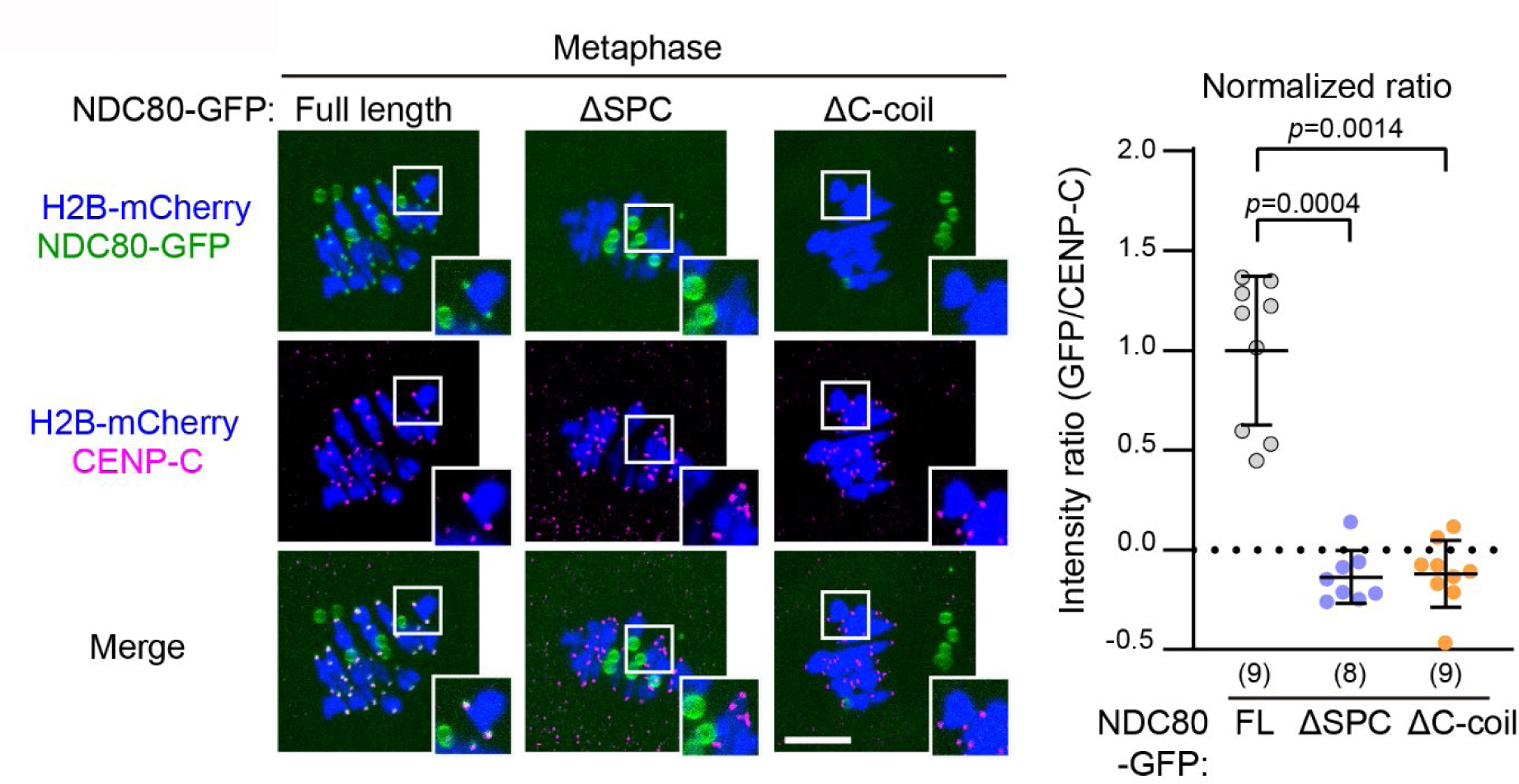
NDC80^ΔSPC^-NUF2 is deficient in kinetochore localization. NDC80^ΔSPC^-GFP is deficient in kinetochore localization. Oocytes were fixed 5 hours after NEBD and immunostained for GFP (NDC80-GFP, green), CENP-C (magenta), and H2B-mCherry (blue). NDC80 (1-642 aa, full length), NDC80^ΔSPC^ (1-611 aa), NDC80^ΔC-coil^ (1-577 aa) were tested. Regions indicated by white boxes are magnified. Z-projection images are shown. The cytoplasm-subtracted intensity of NDC80-GFP signals on kinetochores, relative to that of CENP-C signals, was calculated. Each dot shows the average value of 40 kinetochores in one oocyte. The images and dataset of ‘NDC80-FL-GFP’ were acquired in the same experiments as those in **Fig. S3f** and thus are identical. The number of oocytes from 3 experiments is indicated in parentheses. Scale bar, 10 µm. P-values were calculated with Kruskal-Wallis test with Dunn’s correction.

**Fig. S5:**
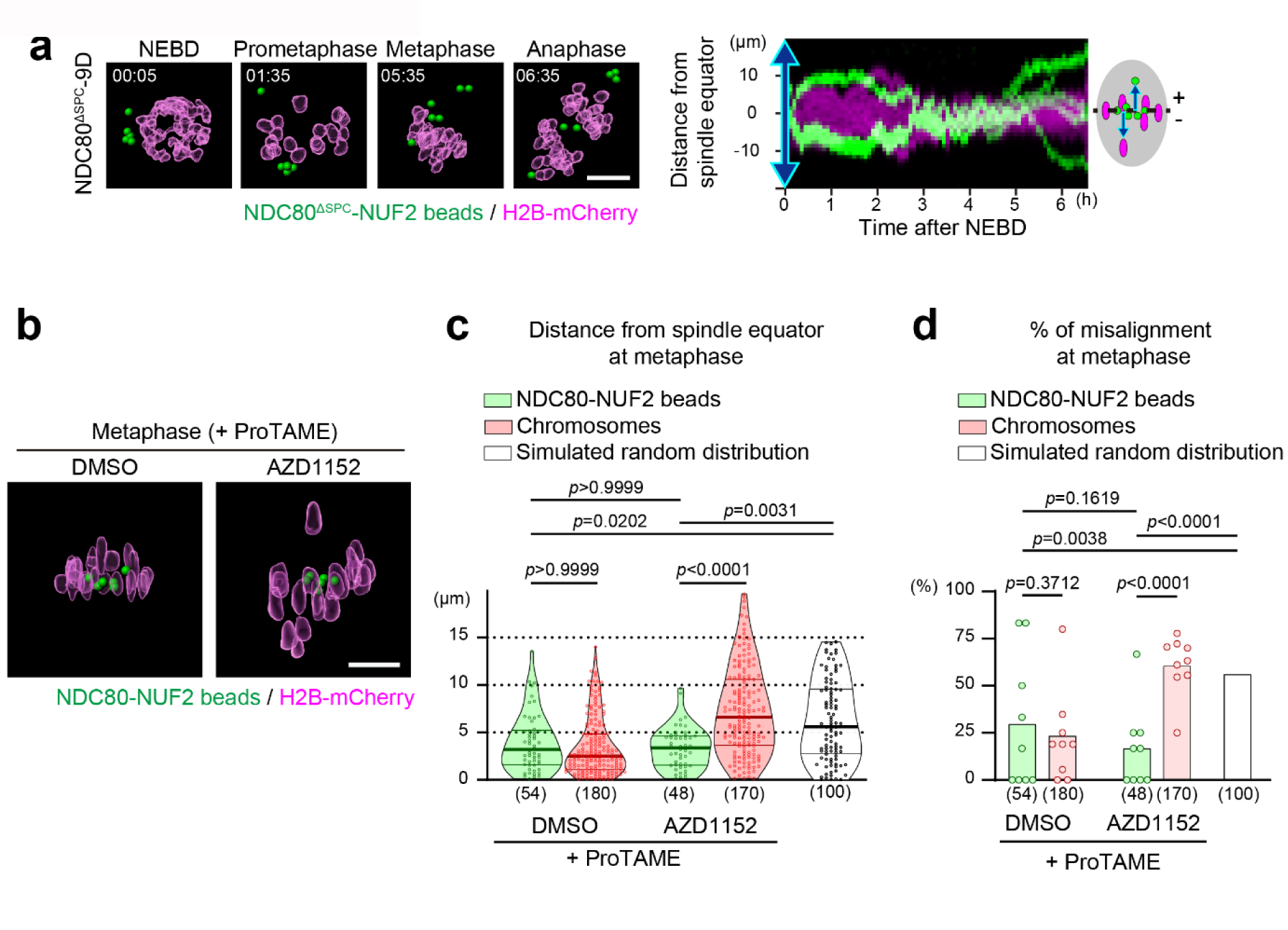
NDC80 phosphoregulation is dispensable for microbead alignment. **(a)** 3D-reconstructed images of chromosomes (H2B-mCherry, magenta) and NDC80^ΔSPC^-9D-NUF2 beads (green). Time after NEBD (hh:mm). Scale bar, 10 µm. The kymograph shows projected signals on the spindle axis over time in chromosomes and NDC80^ΔSPC^-9D-NUF2 beads. The vertical axis shows distance from the spindle equator. For control groups and quantification, see Fig. 4a, b. **(b)** NDC80-NUF2 beads align independently of Aurora B and C. Oocytes carrying NDC80-NUF2 beads were cultured with AZD1152, an inhibitor of Aurora B and C. To prevent precocious anaphase entry due to the inhibition of Aurora B and C, oocytes were additionally treated with ProTAME, an proteasome inhibitor. 3D-reconstructed images of chromosomes (H2B-mCherry, magenta) and NDC80-NUF2 beads (green) at 7 hours after NEBD. Scale bar, 10 µm. **(c)** Alignment efficiency. The distances of NDC80-NUF2 beads from the spindle equator were quantified. The number of beads (from 9 and 9 oocytes, respectively, in 3 independent experiments) is indicated in parentheses. Simulated random distribution of particles within a spindle-like ellipsoid (30 µm in length and 20 µm in width) is used as a reference. P-values were calculated with Kruskal-Wallis test with Dunn’s correction. **(d)** Fractions of misaligned NDC80-NUF2 beads. “Misaligned” refers to positions over 5 µm from the spindle equator at 7 hours after NEBD. Each dot shows the fraction in one oocyte. The number of oocytes (from 3 independent experiments) is indicated in parentheses. P-values were calculated with Fisher’s exact test.

## Notes

### Competing Interest Statement

The authors have declared no competing interest.

## Reference

1. Cheeseman, I. M. The Kinetochore. Cold Spring Harb. Perspect. Biol. 6, a015826 (2014).

2. Musacchio, A. & Desai, A. A Molecular View of Kinetochore Assembly and Function. Biology 6, 5 (2017).

3. Tanaka, T. U. et al. Evidence that the Ipl1-Sli15 (Aurora Kinase-INCENP) Complex Promotes Chromosome Bi-orientation by Altering Kinetochore-Spindle Pole Connections. Cell 108, 317–329 (2002).

4. Liu, D., Vader, G., Vromans, M. J. M., Lampson, M. A. & Lens, S. M. A. Sensing Chromosome Bi-Orientation by Spatial Separation of Aurora B Kinase from Kinetochore Substrates. Science 323, 1350–1353 (2009).

5. Uchida, K. S. K. et al. Kinetochore stretching inactivates the spindle assembly checkpoint. J. Cell Biol. 184, 383–390 (2009).

6. Maresca, T. J. & Salmon, E. D. Intrakinetochore stretch is associated with changes in kinetochore phosphorylation and spindle assembly checkpoint activity. J. Cell Biol. 184, 373–381 (2009).

7. Wan, X. et al. Protein Architecture of the Human Kinetochore Microtubule Attachment Site. Cell 137, 672–684 (2009).

8. Welburn, J. P. I. et al. Aurora B Phosphorylates Spatially Distinct Targets to Differentially Regulate the Kinetochore-Microtubule Interface. Mol. Cell 38, 383–392 (2010).

9. Biggins, S. et al. The conserved protein kinase Ipl1 regulates microtubule binding to kinetochores in budding yeast. Genes Dev. 13, 532–544 (1999).

10. Cheeseman, I. M., Chappie, J. S., Wilson-Kubalek, E. M. & Desai, A. The Conserved KMN Network Constitutes the Core Microtubule-Binding Site of the Kinetochore. Cell 127, 983–997 (2006).

11. DeLuca, J. G. et al. Kinetochore Microtubule Dynamics and Attachment Stability Are Regulated by Hec1. Cell 127, 969–982 (2006).

12. Yue, Z. et al. Deconstructing Survivin: comprehensive genetic analysis of Survivin function by conditional knockout in a vertebrate cell line. J. Cell Biol. 183, 279–296 (2008).

13. Campbell, C. S. & Desai, A. Tension sensing by Aurora B kinase is independent of survivin-based centromere localization. Nature 497, 118–121 (2013).

14. Hengeveld, R. C. C., Vromans, M. J. M., Vleugel, M., Hadders, M. A. & Lens, S. M. A. Inner centromere localization of the CPC maintains centromere cohesion and allows mitotic checkpoint silencing. Nat. Commun. 8, 15542 (2017).

15. Akiyoshi, B. et al. Tension directly stabilizes reconstituted kinetochore-microtubule attachments. Nature 468, 576–579 (2010).

16. Miller, M. P., Asbury, C. L. & Biggins, S. A TOG Protein Confers Tension Sensitivity to Kinetochore-Microtubule Attachments. Cell 165, 1428–1439 (2016).

17. Herman, J. A., Miller, M. P. & Biggins, S. chTOG is a conserved mitotic error correction factor. eLife 9, e61773 (2020).

18. Wei, R. R., Al-Bassam, J. & Harrison, S. C. The Ndc80/HEC1 complex is a contact point for kinetochore-microtubule attachment. Nat. Struct. Mol. Biol. 14, 54–59 (2007).

19. Ciferri, C. et al. Implications for Kinetochore-Microtubule Attachment from the Structure of an Engineered Ndc80 Complex. Cell 133, 427–439 (2008).

20. Espeut, J., Cheerambathur, D. K., Krenning, L., Oegema, K. & Desai, A. Microtubule binding by KNL-1 contributes to spindle checkpoint silencing at the kinetochore. J. Cell Biol. 196, 469–482 (2012).

21. Varma, D. et al. Recruitment of the human Cdt1 replication licensing protein by the loop domain of Hec1 is required for stable kinetochore–microtubule attachment. Nat. Cell Biol. 14, 593–603 (2012).

22. Zhang, G. et al. The Ndc80 internal loop is required for recruitment of the Ska complex to establish end-on microtubule attachment to kinetochores. J. Cell Sci. 125, 3243–3253 (2012).

23. Polley, S. et al. Stable kinetochore-microtubule attachment requires loop-dependent Ndc80-Ndc80 binding. EMBO J. 42, e112504 (2023).

24. Silljé, H. H. W., Nagel, S., Körner, R. & Nigg, E. A. HURP Is a Ran-Importin β-Regulated Protein that Stabilizes Kinetochore Microtubules in the Vicinity of Chromosomes. Curr. Biol. 16, 731–742 (2006).

25. Brunet, S. et al. Kinetochore Fibers Are Not Involved in the Formation of the First Meiotic Spindle in Mouse Oocytes, but Control the Exit from the First Meiotic M Phase. J. Cell Biol. 146, 1–12 (1999).

26. Yoshida, S., Kaido, M. & Kitajima, T. S. Inherent Instability of Correct Kinetochore-Microtubule Attachments during Meiosis I in Oocytes. Dev. Cell 33, 589–602 (2015).

27. Yoshida, S. et al. Prc1-rich kinetochores are required for error-free acentrosomal spindle bipolarization during meiosis I in mouse oocytes. Nat. Commun. 11, 2652 (2020).

28. Veld, P. J. H. in ‘t, Volkov, V. A., Stender, I. D., Musacchio, A. & Dogterom, M. Molecular determinants of the Ska-Ndc80 interaction and their influence on microtubule tracking and force-coupling. eLife 8, e49539 (2019).

29. So, C. et al. A liquid-like spindle domain promotes acentrosomal spindle assembly in mammalian oocytes. Science 364, eaat9557 (2019).

30. Balboula, A. Z. & Schindler, K. Selective Disruption of Aurora C Kinase Reveals Distinct Functions from Aurora B Kinase during Meiosis in Mouse Oocytes. PLoS Genet. 10, e1004194 (2014).

31. Lampson, M. A., Renduchitala, K., Khodjakov, A. & Kapoor, T. M. Correcting improper chromosome–spindle attachments during cell division. Nat. Cell Biol. 6, 232–237 (2004).

32. Rieder, C. L. The Formation, Structure, and Composition of the Mammalian Kinetochore and Kinetochore Fiber. Int. Rev. Cytol. 79, 1–58 (1982).

33. Magidson, V. et al. Adaptive changes in the kinetochore architecture facilitate proper spindle assembly. Nat. Cell Biol. 17, 1134–1144 (2015).

34. Lacefield, S., Lau, D. T. C. & Murray, A. W. Recruiting a microtubule-binding complex to DNA directs chromosome segregation in budding yeast. Nat. Cell Biol. 11, 1116–1120 (2009).

35. Kiermaier, E., Woehrer, S., Peng, Y., Mechtler, K. & Westermann, S. A Dam1-based artificial kinetochore is sufficient to promote chromosome segregation in budding yeast. Nat. Cell Biol. 11, 1109–1115 (2009).

36. Gascoigne, K. E. et al. Induced Ectopic Kinetochore Assembly Bypasses the Requirement for CENP-A Nucleosomes. Cell 145, 410–422 (2011).

37. Sissoko, G. B., Tarasovetc, E. V., Marescal, O., Grishchuk, E. L. & Cheeseman, I. M. Higher-order protein assembly controls kinetochore formation. Nat. Cell Biol. 26, 45–56 (2024).

38. Sarangapani, K. K. & Asbury, C. L. Catch and release: how do kinetochores hook the right microtubules during mitosis? Trends Genet. 30, 150–159 (2014).

39. Regt, A. K. de, Clark, C. J., Asbury, C. L. & Biggins, S. Tension can directly suppress Aurora B kinase-triggered release of kinetochore-microtubule attachments. Nat. Commun. 13, 2152 (2022).

40. Cimini, D. et al. Merotelic Kinetochore Orientation Is a Major Mechanism of Aneuploidy in Mitotic Mammalian Tissue Cells. J. Cell Biol. 153, 517–528 (2001).

41. Hara, M. et al. Centromere/kinetochore is assembled through CENP-C oligomerization. Mol. Cell 83, 2188–2205.e13 (2023).

42. Zhou, K. et al. CENP-N promotes the compaction of centromeric chromatin. Nat. Struct. Mol. Biol. 29, 403–413 (2022).

43. Altemose, N. et al. Complete genomic and epigenetic maps of human centromeres. Science 376, eabl4178 (2022).

44. Magidson, V. et al. The Spatial Arrangement of Chromosomes during Prometaphase Facilitates Spindle Assembly. Cell 146, 555–567 (2011).

45. Heald, R. et al. Self-organization of microtubules into bipolar spindles around artificial chromosomes in Xenopus egg extracts. Nature 382, 420–425 (1996).

46. Politi, A. Z. et al. Quantitative mapping of fluorescently tagged cellular proteins using FCS-calibrated four-dimensional imaging. Nat. Protoc. 13, 1445–1464 (2018).

